# Can food and temperature influence regional connectivity patterns of Bivalvia in fragmented archipelagos? Evidence from biophysical modeling applied to French Polynesia

**DOI:** 10.1101/2023.07.17.549411

**Authors:** H. Raapoto, C.J. Monaco, S. Van Wynsberge, R. Le Gendre, J. Le Luyer

## Abstract

Larval dispersal and connectivity are key processes that drive marine metapopulation dynamics, and therefore should be well characterized when designing effective management strategies. While temperature and food availability can structure marine species connectivity patterns, their relative contribution has not been thoroughly investigated in highly fragmented archipelagos. We used biophysical modeling of larval dispersal to explore the connectivity patterns of species with complex life-cycles across French Polynesia (FP), a territory formed by more than a hundred small, geographically isolated islands covering an area as large as Europe. We first simulated ten years of larval dispersal to investigate the spatial and temporal (seasonal and interannual) variability in larval dispersal pathways for different hypothetical species exhibiting a range of Larval Precompetency Period (LPP) values. Then, using the black-lip pearl oyster (*Pinctada margaritifera*) as a model species, we accounted for variability in the LPP induced by temperature and food availability, as derived from a Dynamic Energy Budget (DEB) model. The model showed that food availability and meso-scale turbulence (eddies) in the Marquesas jointly constrained larval dispersal, reducing its potential connectivity with other archipelagos in FP. However, accounting for food and temperature effects on larval development, barely changed the connectivity pattern at regional scale due to the remoteness of this archipelago. The DEB simulations further revealed seasonal and interannual variability in connectivity driven by environmental conditions. Our results highlight the importance of considering temperature and food in biophysical models to adequately capture dispersal, connectivity and to identify appropriate management units at the regional scale.

## 1 Introduction

Demographic connectivity as defined by the exchange of individuals among geographically distinct populations, is vital for the persistence of metapopulations (Williams & Hastings, 2013), for the recovery of depleted populations following perturbations (Lipcius et al., 2008), and for maintaining genetic diversity (Trakhtenbrot et al., 2005). Characterizing larval dispersal and connectivity patterns of marine species is therefore crucial for quantifying population dynamics and to support effective resource-management policies (Cowen et al., 2007; Kritzer & Sale, 2004; Lipcius et al., 2008). However, assessing connectivity in marine systems remains a challenging task because most marine species have complex life histories involving planktonic larval stages.

Various methods exist to infer marine population connectivity, including genetic approaches (Lowe & Allendorf, 2010), geochemical markers (e.g. microchemical signatures in shells), *in situ* larval sampling techniques, or larval dispersal through hydrodynamic or biophysical modelling (see review in <u>Jahnke & Jonsson, 2022)</u>. While empirical methods are often inadequate for species occurring across large geographic ranges, especially hyper fragmented archipelagos, biophysical models emerge as suitable because they can account for both large spatial and temporal scales (Dickey & Bidigare, 2005; Legrand et al., 2022; Nichols & Raghukumar, 2020) . Lagrangian models in particular, which allow tracing the displacement of individual particles, are increasingly used for a wide range of applications (van Sebille et al., 2018) including describing the trajectory of marine debris (Dobler et al., 2019; Maes & Blanke, 2015), guiding search and rescue operations (Durgadoo et al., 2019; Hart-Davis & Backeberg, 2021), estimating the age of water masses (Deleersnijder et al., 2001), and mapping the dispersion (Wood et al., 2016) and connectivity of marine larvae (Quinn et al., 2017; Wolanski et al., 2021).

Following Lagrangian models, the population connectivity of marine organisms with planktonic larval stage can be formalized as the probability of transport between two sites. This depends mainly on (i) the speed and direction of transport, as determined by oceanographic currents for planktonic larvae, and (ii) the Pelagic Larval Duration (PLD; i.e., time spent drifting before settlement). The PLD varies substantially across species and environmental conditions (from few hours to >100 days; <u>O’Connor et al., 2007)</u>. The lower limit of a species’ PLD is determined by the Larval Pre-competency Period (LPP; also known as settlement competency period), defined as the minimum time required for the larva to be able to settle. Once LPP is reached, larvae can fix as long as they reach adequate habitat conditions <u>(O’Connor et al., 2009; Shima & Swearer, 2009; Wilson & Harrison, 1998)</u>. LPP is genetically controlled (species-specific) but it is also a plastic trait that varies considerably in response to environmental conditions, which can have important consequences for larval connectivity. For example, if warm temperatures accelerate larval development, the LPP and larval dispersal are reduced and connectivity patterns can be altered (O’Connor et al., 2007; Raventos et al., 2021). While the importance of temperature variability is generally acknowledged in larval connectivity models, few studies consider the role of food availability in larval development and growth (Sangare et al., 2020a). And even though LPP or PLD are often included in biophysical models of dispersion, values are typically fixed based on empirical observations in controlled environments, thus rarely considering the dynamic environment encountered by larvae in the wild. An elegant and increasingly popular approach to account explicitly for the influence of temperature and food availability on larval dispersal is to combine biophysical and bioenergetic models, which is now possible thanks to recent advances in the fields of physical oceanography and physiological ecology (Falcini et al., 2020). Dynamic Energy Budget (DEB) models (Kooijman, 2010) provide powerful means to estimate LPP based on the local temperature and food conditions experienced by individual larvae, thus potentially improving the predictions from biophysical dispersion models (Sangare et al., 2019, 2020a).

The French Polynesian (FP) territory provides an interesting playground to investigate larval dispersal and population connectivity patterns because of its sheer size (comparable to that of Europe) and the presence of numerous islands organized in five highly-fragmented archipelagos that provide a diversity of environmental conditions. Even though the tropical ocean is typically characterized by a low productivity, higher temperature, and a weaker seasonal signal than temperate latitudes (Claustre et al., 2008; Dufour et al., 1999), large inter archipelago variation exist in FP, notably between the Marquesas and the Gambier. Furthermore, ocean circulation and environmental conditions are influenced on a seasonal scale but also on an interannual scale. Climate patterns such as ENSO (El Niño - Southern Oscillation) are known to impact the physico-chemistry of the Pacific Ocean (Delcroix, 1998; Gouriou & Delcroix, 2002; McPhaden & Zhang, 2004). Thus, such forcing might affect larval dispersal and connectivity in the area. Few studies, however, have aimed to quantify larval dispersal and connectivity in the region (but see <u>Martinez et al., 2007</u> for macroalgae drifts). While intra-lagoon larval dispersal has been examined for some FP atolls by either coupling hydrodynamic and bioenergetic models or by combining hydrodynamic and genetics information (Reisser et al., 2019), this has not been done across the whole of FP.

A quantitative understanding of the hydrodynamic and biological forces driving population connectivity across FP is of particular interest when considering its various bivalve species (Tröndlé & Boutet, 2009). These mollusks play significant roles in the local marine ecosystems, particularly in the benthic habitats. They are instrumental in the cycling of nutrients and contribute to the overall productivity of the marine environment (Vaughn & Hoellein, 2018). Among the many species of bivalves present in FP, the black-lip pearl oyster (*Pinctada margaritifera*) is a species of high cultural and economic value that supports the local pearl industry (Andréfouët et al., 2012). *Pinctada margaritifera* has been the focus of numerous research projects, and significant contributions on population genetics provide insights into the potential degree of genetic structuring at basin scale (Lal et al., 2017) and also within and across FP atolls (Arnaud-Haond et al., 2004, 2008; Lemer & Planes, 2014; Reisser et al., 2019). This work, however, is limited because even though it can describe the existing genetic structure of populations to infer connectivity, it is unable to unravel the ecological processes that produce them.

In this study, we first use biophysical modeling and graph-theory to explore overall larval dispersal patterns in FP for different LPP values that are relevant for a wide range of species. We run these simulations over ten years to specifically investigate the spatial extent, the seasonal influence, and the interannual variability of larvae trajectories. Then, using *P. margaritifera* as model species, we computed the same dispersal simulations but using LPP as estimated from an along-track temperature- and food-dependent DEB model. This coupled biophysical-bioenergetics model allows, for the first time, exploring the role of temperature and food availability on the connectivity patterns of sessile broadcast-spawner species across FP islands.

## 2 Methods

### 2.1 Study area

FP, located in the tropical South Pacific (155°W-130°W, 5°S-30°S; Fig. 1), is a vast territory spread over an area as large as Europe (∼4.5 million km^2^). Organized in 5 archipelagos with a total of 118 volcanic islands and atolls (Duncan & McDougall, 1976), only a small fraction of FP sits above sea level (4,200 km², <0.1 % of total area). Islands and atolls display extraordinary geomorphological variation, ranging in size between <1 and ∼2,000 km^2^ (Andréfouët et al., 2001; Galzin et al., 1994; Rougerie et al., 1997). While most of the Polynesian atolls and islands have a lagoon, high islands forming the northernmost archipelago (Marquesas) do not because of their drowned reefs (Rougerie et al., 1997). For islands and atolls that have lagoons, the degree of connectedness between the lagoon and ocean vary considerably from open to close, depending on the presence, number, and characteristics of reef passes and of shallow channels that cross the atoll rim (called *hoa*).

**Figure 1:**
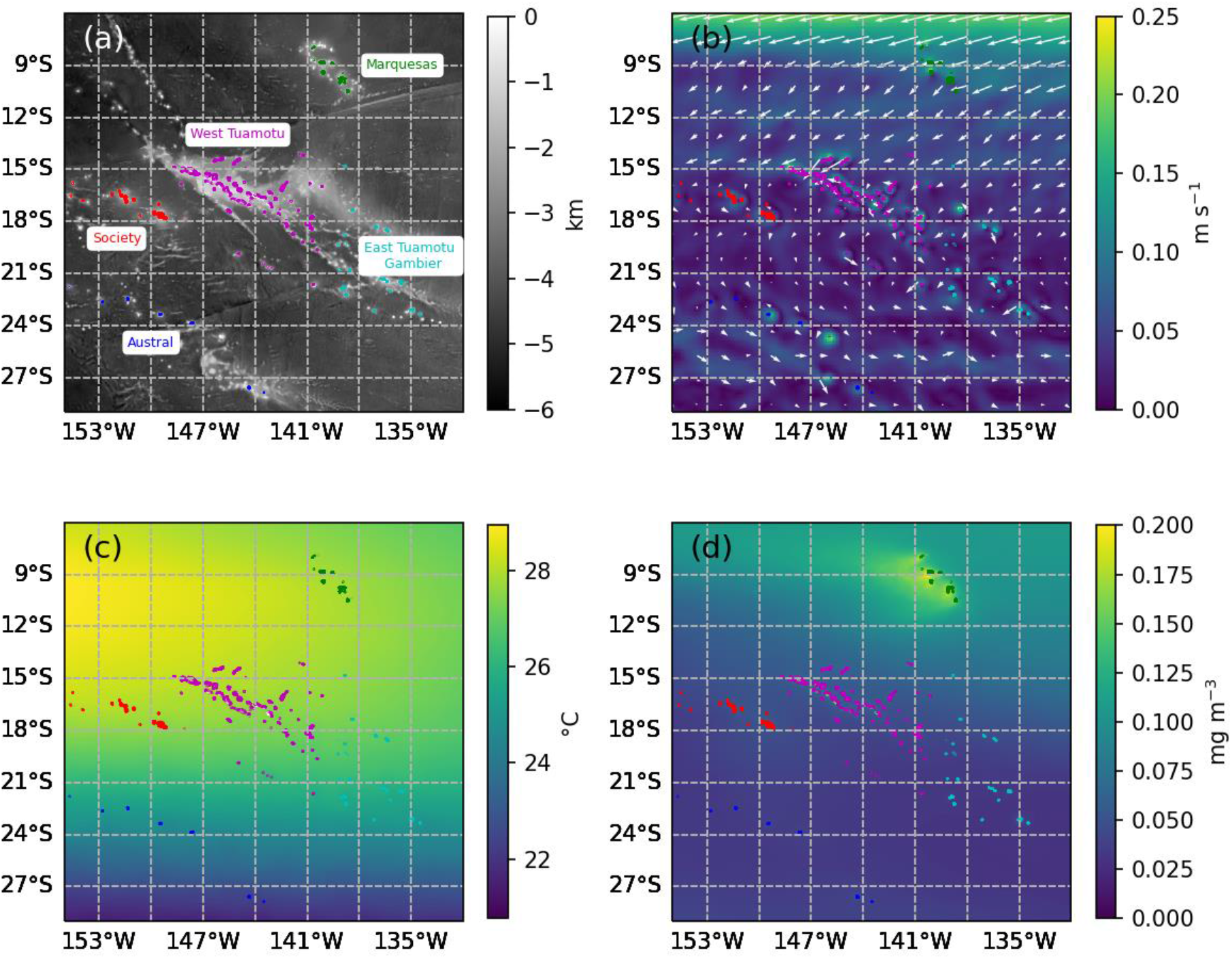
a) Bottom topography of FP with the Marquesas (green), Western Tuamotu (pink), Society (red), Eastern Tuamotu – Gambier (cyan), and Austral (blue) archipelagos. (b) Mean current speed (b) and (c) temperature averaged over 2010 to 2019 across the first 50 m depth from Mercator Ocean global reanalysis (GLORYS12V1). (d) Mean surface chlorophyll-a averaged on the same time period from Global Ocean Colour (GlobColour) data.

### 2.2 Biophysical modeling and ocean data

To build the biophysical model for FP, we used current velocity and temperature data from the Mercator Ocean global ocean reanalysis (GLORYS12V1) product, which provides daily data (interpolated to a higher temporal resolution for the Lagrangian experiment – see section 2.3 *Lagrangian particle tracking*) on a spatial horizontal resolution of 1/12° (∼9 km) and across 50 vertical levels. This product is available through the Copernicus Marine Environment Monitoring Service (CMEMS, https://marine.copernicus.eu/) described by <u>Lellouche et al. (2018)</u> . For this study, we averaged current velocities and temperatures across the first 50 meters to match the vertical distribution of *P. margaritifera* larvae (Thomas, Garen, et al., 2012).. This also provide a more general picture of the flow field and thermal conditions that larval particles are likely to experience. Chlorophyll-*a* concentration data (Chl-a), a proxy for phytoplankton biomass (i.e., food availability for the larvae) were extracted from the product Copernicus-GlobColour based on multi-satellite observations, which provides daily estimates with a 4-km resolution (Maritorena et al., 2010). All data used in this study were extracted for the 2010-2019 period.

### 2.3 Lagrangian particle tracking

To model the Lagrangian drift of larvae we used the software Ocean Parcels (Delandmeter & van Sebille, 2019; Lange & van Sebille, 2017). Ocean Parcels has been used in a variety of oceanographic applications (Escalle et al., 2019; Onink et al., 2019; Scutt Phillips et al., 2018; Van Sebille et al., 2019), and was parameterized in this study as follows:

#### Particle emission

For the whole study period (2010-2019), 100 particles were released daily from each of the 102 biggest islands of FP. Islands that were in close proximity (< 2 km; i.e., Ravahere-Marokau) or those with a shared lagoon (i.e., Mangareva, Raiatea-Tahaa) were considered as one unique island. Particle initial positions were generated randomly from a 2-km buffer area extending between the crest of the outer reef and the reef slope. The locations of reef geomorphological units were extracted from the Millennium Coral Reef Mapping Product (MCRMP; <u>Andréfouët et al., 2006)</u>. A total of 10,200 particles per day were released, representing a total of 37,250,400 particles for the 10 years of simulation (i.e., 365,200 per island).

#### Particle drift

In Ocean Parcels we set the time step of iteration to 1 hour and the position of larvae was recorded every 3 hours, which is lower than the ratio of cell size (9 km) to maximum current velocity in the area (∼20 cm s^-1^). This prevents particles to cross more than one boundary or hydrodynamic cell in a single time step and thus, prevents missing positions that might reach the small islands. Advection was simulated using a fourth-order Runge-Kutta integration scheme. Current velocity at the particle location was obtained through spatial and temporal linear interpolation of the flow field data. Because the resolution of the hydrodynamic forcing does not properly resolve dispersion related to turbulent eddies, a stochastic ‘random walk’ impulse was applied to each modelled larva (Quinn et al., 2017; van Sebille et al., 2018). Temperature and Chl-a at each particle location were obtained through spatial and temporal linear interpolation and recorded every 3 hours.

Once LPP was reached (see section 2.4 *Larval dispersal scenarios tested*), we allowed particles to settle as soon as they crossed a patch of suitable habitat (i.e., hard substrate surrounding an island or atoll). Because most FP islands are too small to be properly captured by the resolution of GLORYS data, masks of hard substrates were designed on the basis of MCRMP and used to determine when particles reached suitable habitat. Larvae were considered as settled when they entered a 2-km buffer zone around these masks (same as defined for the initial positions). We tracked particles for a maximum of 35 days, as this encompasses the PLD for many tropical marine species (O’Connor et al., 2007; Wellington & Victor, 1989), including *P. margaritifera* (Sangare et al., 2019; Thomas et al., 2014b). Beyond this period of 35 days, particles that did not encounter a suitable habitat were considered lost.

### 2.4 Larval dispersal scenarios tested

Nine scenarios were considered to investigate the relative importance of LPP and environmental conditions in the global connectivity patterns. A baseline scenario (scenario 0) forced larvae to drift until the end of the simulation (35 days), to assess the maximal extent of the dispersal kernel that can be expected considering currents velocity and direction in FP.

A set of 7 scenarios included connectivity matrices for a wide range of marine species exhibiting a spectrum of LPPs (and maximum PLD; PLD_max_), from 3 to 23 days (Table 1). The LPP was fixed to two thirds of the PLD_max_ in our scenarios to consider the fact that the longer the pre-competent period, the greater the capacity for delaying metamorphosis (Pechenik, 1990; Wellington & Victor, 1989).

**Table 1:** larval pre-competency period (LPP) and maximum pelagic larval duration (PLDmax) for the eight scenarios considered in this study

| Scenario | Settlement parametrization | LPP (days) | PLD<sub>max</sub> (days) |
| --- | --- | --- | --- |
| 0 | No-settlement scenario | - | 35 |
| 1 | Settlement as soon as crossing suitable habitat when LPP reached | 3 | 5 |
| 2 |  | 7 | 10 |
| 3 |  | 10 | 15 |
| 4 |  | 13 | 20 |
| 5 |  | 17 | 25 |
| 6 |  | 20 | 30 |
| 7 |  | 23 | 35 |
| 8 |  | From DEB | 35 |

The eighth scenario specifically considered the case study of *P. margaritifera*. We determined the LPP of each simulated larva based on predictions from a Dynamic Energy Budget model for the species <u>(Sangare et al., 2020;</u> see section *2.5.* *Spatial DEB modeling to determine LPD*). In all scenarios, no mortality curve nor predation were simulated.

### 2.5 Spatial DEB modeling to determine LPP

To determine the effective pre-competency period of each individual larva of *P. margaritifera* released in our simulations (scenario 8), we estimated growth and development using dynamic energy budget theory (Kooijman, 2010). Based on a robust theoretical framework that is consistent with and can explain long-standing empirical models in biology (e.g., von Bertalanffy growth) (Sousa et al., 2010), DEB theory allows quantifying the energy use and allocation of organisms experiencing variable environmental conditions (temperature and food availability). Throughout the life of an organism, DEB models can explicitly predict key life-history traits including growth, reproduction, and notably maturity level (Monaco et al., 2014). Here we used the DEB model described for *P. margaritifera* by <u>Sangare et al., 2019,</u> <u>2020</u>; parameter values available at https://www.bio.vu.nl/thb/deb/deblab/add_my_pet/entries_web/Pinctada_margaritifera/Pincta da_ margaritifera_res.html), which has been calibrated and validated using data on larva and adult life stages (Fournier, 2011; Thomas et al., 2011). The model considers that larvae grow exponentially until reaching a maturity threshold that marks the onset of metamorphosis, thus defining the end of the LPP. We began each simulation with new-born larvae that then drifted in the water column following the modelled Lagrangian trajectories. At each daily time step, we estimated the larva size and maturity depending on the sea surface temperature and Chl-a data. Since our DEB model does not allow a proper definition of PLD_max_, the same previous statement was used for linking LPP and PLD_max_ (PLD_max_ defined as three halves the LPP). Once particles reached this value, they were considered lost.

### 2.6 Data analysis

#### Dispersal and connectivity metrics

First, the probability of transport from an island or group of islands *i* to an island or group of islands *j* (*P_i→j_*) was calculated as the proportion of larvae emitted from the island *i* that settled at *j*. Then, four larval dispersal metrics (Lett et al., 2015) were calculated based on *P_i→j_*, at the scale of the islands. The settlement success for island *i* (SS_i_) was calculated as the proportion of particles emitted from *i* that could settle (either in the same island/group of islands or elsewhere; eq. 1), while the recruitment success (RS_i_) for an island *i*, is the ratio of the number of particles settled on *i* and the total number of particles emitted from the whole dataset on the time period considered (eq. 2). Local retention at island *i* (LR_i_) is defined by *P_i→i_*, and self-recruitment (SR_i_) is the ratio of locally produced settlement to settlement of all origins at a site (eq. 3).

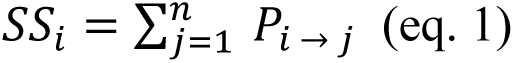

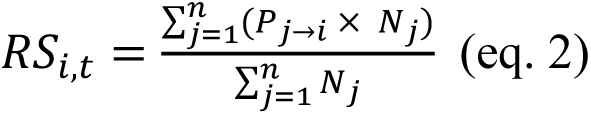

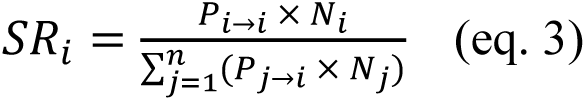

With *N_i_ and N_j_* the number of larvae emitted from island *i* and *j,* respectively.

#### Seasonal and interannual trends

A seasonality analysis was performed on the total RS from all islands to determine the monthly recruitments. They were obtained by computing the average and standard deviation on monthly data over the 10 years of simulation. To visualize interannual dispersal trends, monthly recruitments (RS) were filtered with the STL (Seasonal Trend decomposition procedure based on Loess) additive scheme (Cleveland et al., 1990). This filtering procedure decomposes a time series into trends, seasonal and remaining components, which is particularly appropriate for extracting the interannual and trend signals from non stationary and noisy climate datasets (Terray, 2011). The STL procedure decomposes the analyzed X(t) monthly time series into three terms:

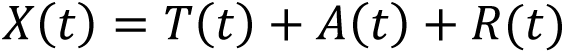

The T(t) term quantifies the trend and low-frequency variations in the time series. The A(t) term describes the annual cycle and its modulation through time. Finally, R(t) is the residual term containing the interannual signal and the noise present in the data. A comparison of RS trends and the Southern Oscillation Index (SOI) was also performed to assess the possible influence of ENSO on larval dispersion. The SOI data were retrieved from the NOAA dataset (https://www.cpc.ncep.noaa.gov/data/indices/soi).

#### Network analysis

We used a network analysis based on graph theory (Koutrouli et al., 2020) on the probability of transport from an island *i* to an island *j* (*P_i→j_*) to i) explore connectivity patterns across islands over the 10-year study period (2010-2019), ii) identify putative island clusters based on *degree* metrics and iii) compare the clustering across the 8 scenarios tested. Briefly, each node in our network corresponds to islands or atolls in FP (n = 102; supp. Table 1) while directed edges (*from*, *to*) represent mean dispersal rates across nodes. The overall network connectivity was estimated, and common graph metrics were assessed for each node to identify atolls or islands of priority for resource management purposes. First, we used the *network density* (ratio of observed edges to the number of possible edges in the network) as a proxy of overall network fragmentation. We used the node’s *degree* (the number of connections ingoing and outgoing from a node) as a measure of the importance of the node to act as source or sink. Finally, we explored sub-structuring (i.e., clusters) based on the hierarchical Louvain’s method (Blondel et al., 2008). All network analyses were conducted with the *scikit-network* python library (Bonald et al., 2020). Dispersal metrics (SS, RS, LR and SR) previously computed at the scale of the islands were also computed at the scale of the clusters i.e group of islands) and the archipelagos. For testing differences across clusters or islands, we used yearly values and non-parametrical Kruskal-Wallis rank sum tests. Pairwise comparison tests were then computed using Wilcoxon rank sum exact tests and Bonferroni correction.

## 3 Results

### 3.1 Spatial and temporal patterns of temperature, food, and currents

The surface ocean circulation mainly followed the south pacific gyre over the study period. In the northern part of the area, the South Equatorial Current (SEC) flowing westward/southwestward was the main current affecting FP and was stronger around the Marquesas archipelago with yearly average velocities around 20 cm s^-1^, compared to 10 cm s^-1^ in the rest of the area (Figure 1b).

The yearly mean ocean temperature averaged across the upper 50 meters ranged between 22°C in the Austral archipelago and 28°C in the western Tuamotu and Society islands (Figure 1c). In the north-east area, the imprint of the equatorial upwelling induces slightly colder waters (< 27.5 °C). While in the north-west, the south-eastward imprint of the warm pool induces slightly warmer water (> 28 °C).

Oceanic water masses in FP are nutrient-depleted with low Chl-a concentrations (Figure 1d). The mean surface Chl-a shows a south-west/north-east gradient due to the contribution of nutrient-rich waters from the equatorial upwelling. The highest values are observed around the Marquesas archipelago.

### 3.2 Potential particle dispersal over 35 days (Scenario 0)

For the 10 years of simulation, the spatial extent of particles that drifted for 35 days ranged from 0 to 1463 km (scenario 0; Figure 2). This scenario predicted high isolation of the Austral and Marquesas archipelagos, but high connections between the other archipelagos. In this scenario, relatively few particles emitted from the Society archipelago could reach the Austral and West Tuamotu archipelagos (Figure 2b), and none could reach the Marquesas and East Tuamotu-Gambier archipelagos. Particles from the West Tuamotu islands dispersed randomly, albeit a slight south-westward shift of the maximum distribution (Fig. 2c), presumably associated to the main current (Fig. 1b). West Tuamotu particles reach most of the islands from the Society and East Tuamotu-Gambier archipelagos. Particles from the East Tuamotu-Gambier archipelago spread northwestward but could reach the Tuamotu West only (Figure 2d). Larvae released from the Marquesas archipelago, which is the most affected by the SEC (Figure 1b), tend to spread south-westward. However, they could hardly reach the West Tuamotu in 35 days (Figure 2e). Similarly, particles emitted from the Austral could not reach the other archipelagos in 35 days (Figure 2f).

**Figure 2:**
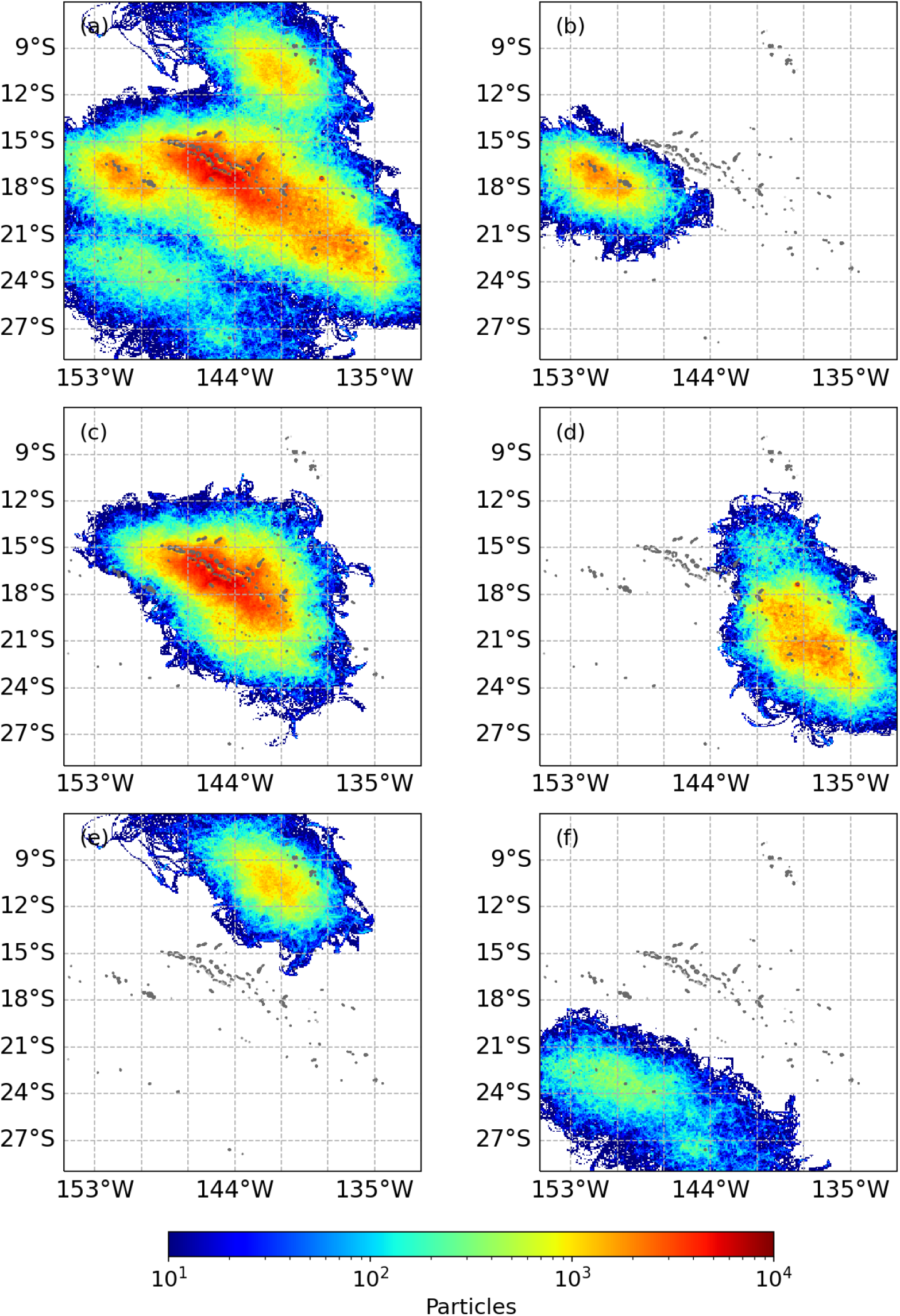
Modelled dispersal paths after a duration of 35 days without settlement (scenario 0), expressed as larval densities and calculated as the number of particles in each cell on a 1/100° grid for (a) all FP, (b) larvae emitted from the Society only, (c) from the West Tuamotu only, (d) from the East Tuamotu-Gambier, (e) from the Marquesas, and (f) from the Austral archipelagos. Note that in this scenario the positions of larvae exclude mortality or settlement. Animation for FP can be viewed in the Supplementary Materials section.

### 3.3 Potential connectivity across a range of LPP scenarios (scenarios 1 to 7)

After ten years of simulation, out of the 37,250,400 particles generated all over FP, only 2.10 and 8.04 % larvae reached a suitable substrate for LPP scenarios of 3 and 23 days, respectively (Table 2). Mean total recruitment success (RS) and mean distance traveled increased linearly with LPP (Table 2).

**Table 2:** Total mean settlement success (SS), recruitment success (RS), self-recruitment (SR), local retention (LR), distance of settlement to source, correlation between SS and SOI and number of clusters over the 10 years of simulation for the various scenarios. Letters indicate significance at Bonferroni adjusted P < 0.05. The number of clusters is computed on 10 years of simulation.

| Scenario | LPP | SS (%) | RS (%) | SR (%) | LR (%) | Distance of settlement to source (km) | Correlation between SOI and SS | p-value | Number of clusters |
| --- | --- | --- | --- | --- | --- | --- | --- | --- | --- |
| 1 | 3 | 2.10 ± 0.16<sup>e</sup> | 0.02 ± 0.00<sup>e</sup> | 63.30 ± 3.17<sup>f</sup> | 1.22 ± 0.14<sup>b</sup> | 27.83 ± 2.57<sup>f</sup> | -0.55 | 5.15 10<sup>-11</sup> | 42 |
| 2 | 7 | 3.04 ± 0.15<sup>f</sup> | 0.03 ± 0.00<sup>f</sup> | 48.82 ± 3.07<sup>g</sup> | 1.39 ± 0.15<sup>bc</sup> | 59.47 ± 5.16<sup>g</sup> | -0.44 | 3.98 10<sup>-7</sup> | 25 |
| 3 | 10 | 4.86 ± 0.27<sup>a</sup> | 0.05 ± 0.00<sup>a</sup> | 40.22 ± 2.57<sup>a</sup> | 1.72 ± 0.20<sup>a</sup> | 89.89 ± 7.03<sup>a</sup> | -0.60 | 7.07 10<sup>-13</sup> | 19 |
| 4 | 13 | 6.18 ± 0.35<sup>b</sup> | 0.06 ± 0.00<sup>b</sup> | 34.51 ± 2.56<sup>b</sup> | 1.74 ± 0.21<sup>a</sup> | 122.04 ± 9.95<sup>b</sup> | -0.61 | 1.31 10<sup>-13</sup> | 14 |
| 5 | 17 | 6.28 ± 0.45<sup>bc</sup> | 0.06 ± 0.00<sup>bc</sup> | 29.86 ± 1.87<sup>c</sup> | 1.47 ± 0.20<sup>abc</sup> | 157.67 ± 13.02<sup>c</sup> | -0.49 | 9.54 10<sup>-9</sup> | 12 |
| 6 | 20 | 7.25 ± 0.57<sup>cd</sup> | 0.07 ± 0.00<sup>cd</sup> | 26.53 ± 1.70<sup>d</sup> | 1.42 ± 0.20<sup>abc</sup> | 188.78 ± 15.13<sup>d</sup> | -0.41 | 2.71 10<sup>-6</sup> | 11 |
| 7 | 23 | 8.04 ± 0.64<sup>d</sup> | 0.08 ± 0.01<sup>d</sup> | 23.70 ± 1.02<sup>e</sup> | 1.36 ± 0.18<sup>bc</sup> | 220.04 ± 17.99<sup>e</sup> | -0.38 | 1.49 10<sup>-5</sup> | 10 |
| 8 | DEB | 8.65 ± 0.99<sup>d</sup> | 0.08 ± 0.01<sup>d</sup> | 25.35 ± 1.67<sup>de</sup> | 1.63 ± 0.28<sup>ac</sup> | 193.34 ± 1.68<sup>d</sup> | -0.21 | 0.02 | 12 |

Monthly settlements revealed no apparent seasonal trend emerging from the different scenarios 1 to 7 (Figure 3a). The settlement trend obtained through the STL analysis is shown in Figure 3b (see Supp Mat for the observed, seasonal, and residual signals). All scenarios with a fixed LPP show the lowest SS in 2011, when the SOI is highly positive (La Niña), and the highest in 2014, when the SOI is becoming negative (El Niño). Indeed, both signals from settlements and SOI present a weak negative correlation for all seven scenarios (Table 2). Low SS trend was also found in 2011 for scenario 8 (DEB), but the lowest SS was obtained in 2017 for this scenario, when the SOI was moderately positive (La Niña).

**Figure 3:**
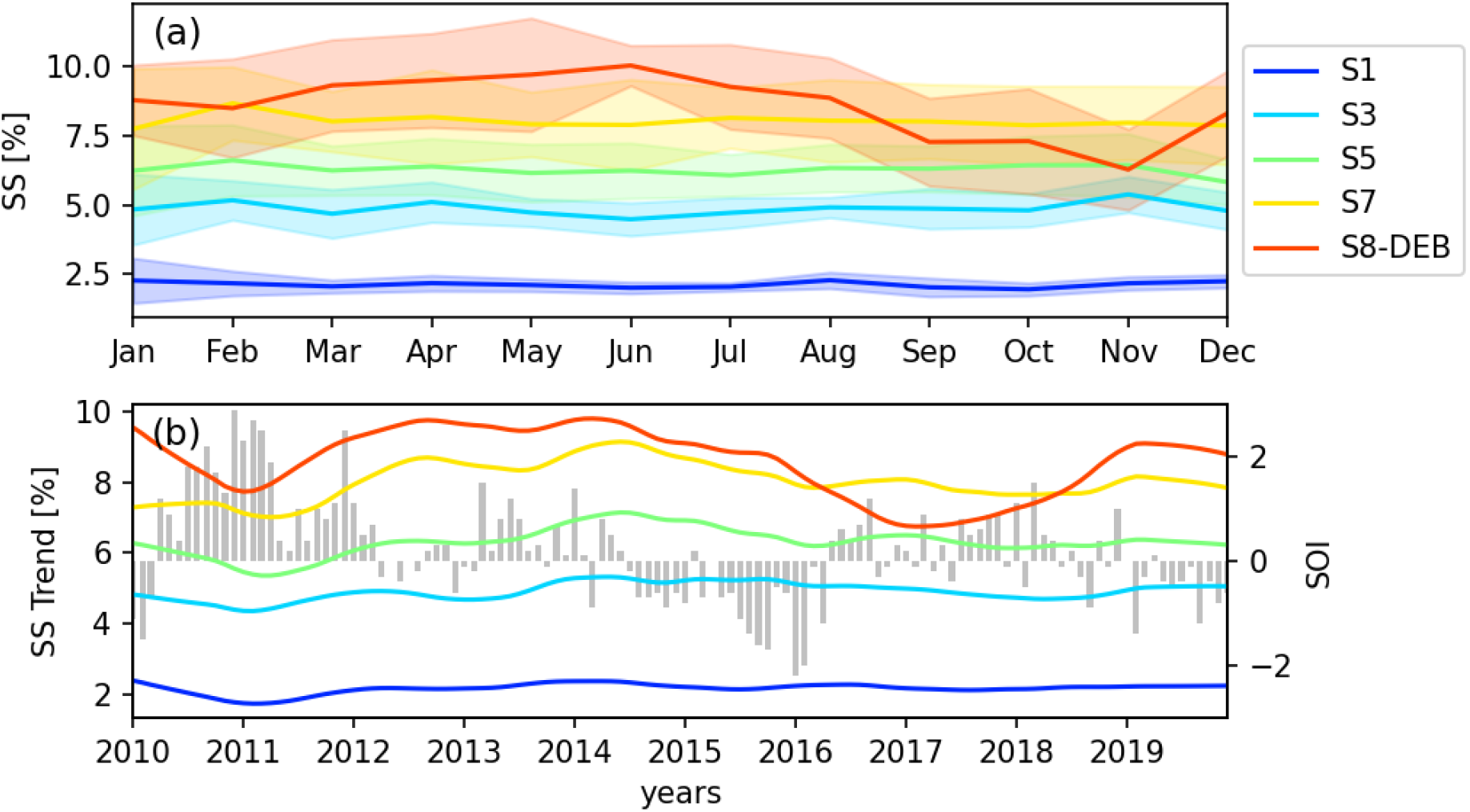
(a) Time series of the monthly average SS, with shades representing their standard deviations and (b) the interannual SS trend extracted from the STL over the full dataset (all archipelagos included). For ease of representation, only scenarios 1, 3, 5, 7 and 8 are represented in blue, light blue, green, yellow and orange, respectively. The SOI is represented in gray bar plot for the same period.

Potential connectivity matrices for LPP of 3, 10, 17 and 23 days, as well as the DEB model-derived LPP, are shown inFigure 4 Figure 4. For the shortest LPP (3 days), apart SR, only few connections were possible with neighboring islands (Table 2 and Figure 4a). Indeed, the mean traveled distance for these particles was around 28 km (Table 2). Therefore, only very close group of islands might share larvae such as those in the Western Tuamotu. With longer LPP, potential connectivity matrices were more complex with connections appearing between more distant islands and exports being possible between different archipelagos (Figure 4b, c, and d).

**Figure 4:**
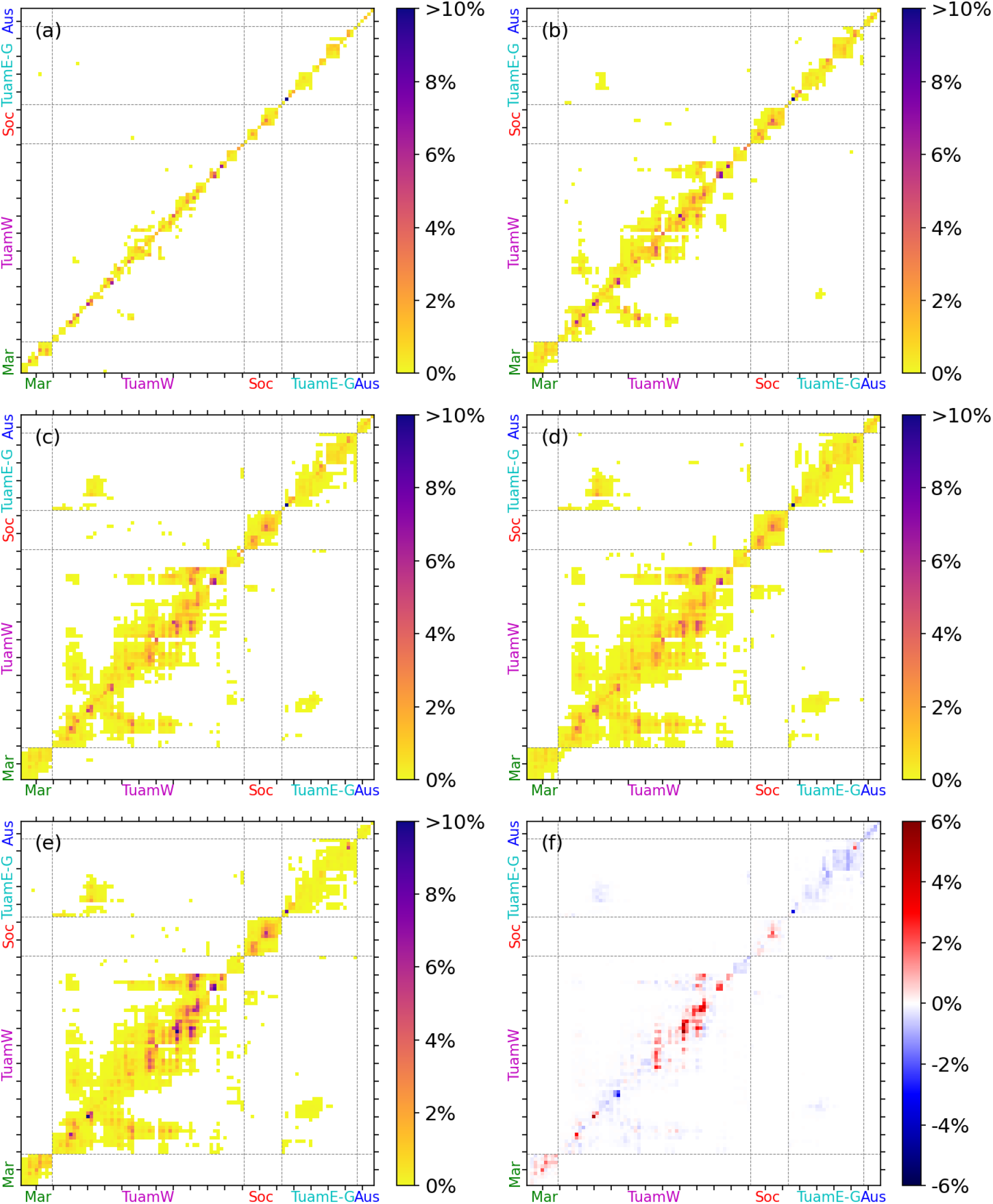
Potential connectivity matrices for (a) scenario 1 (LPP of 3 days), (b) scenario 3 (LPP of 10 days), (c) scenario 5 (LPP of 17 days), (d) scenario 7 (LPP of 23 days) and (e) scenario 8 (LPP based on the DEB model). The difference between scenario 8 and 7 is shown in (f). Source islands are on the left axis and destination islands on the bottom one. Indexes from the Marquesas (Mar), Western Tuamotu (TuamW), Society (Soc), Eastern Tuamotu - Gambier (TuamE-G), and Austral (Aus) archipelagos are colored in green, pink, red, cyan, and blue, respectively. The islands in the matrices are sorted according to the main ocean currents (see Supp Table 1)

For the 10 years of simulation, network fragmentation decreased with increasing LPP as shown by the reduction of the number of clusters and linear increase in network density (Fig. 5a). Years 2011 (La Niña) and 2013/2014 had the minimum and maximum network density, respectively (Figure 5a).

**Figure 5:**
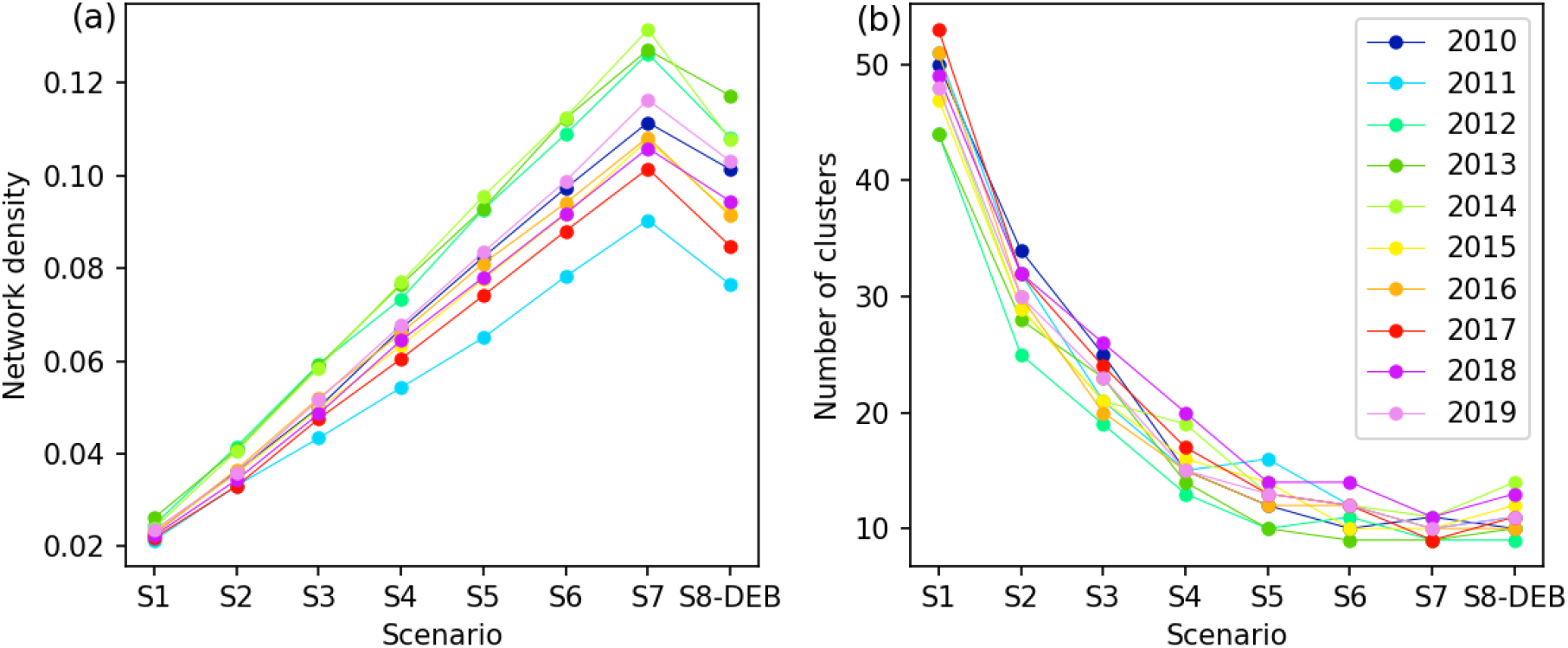
(a) Network density and (b) number of clusters per year and according to the different scenarios. Points were joined with lines for better visualization.

Islands and atolls grouped into clusters are shown in Figure 6 and the total number of clusters identified for each scenario over the 10 years in Table 2. For 3-day LPP (i.e., 5-day maximum PLD; scenario 1), islands were grouped in 42 clusters with the largest one formed by 11 (out of 102) islands (Figure 6a). As LPP increased, the clusters were joined together to form larger components, increasing the overall connectivity in FP. At the longest PLD_max_ (35 days), islands were grouped into 10 clusters, with the largest one formed by 28 islands (27.4% - black in Figure 6d). The region being so fragmented, none of the configurations presented here allowed the system to be fully connected.

**Figure 6:**
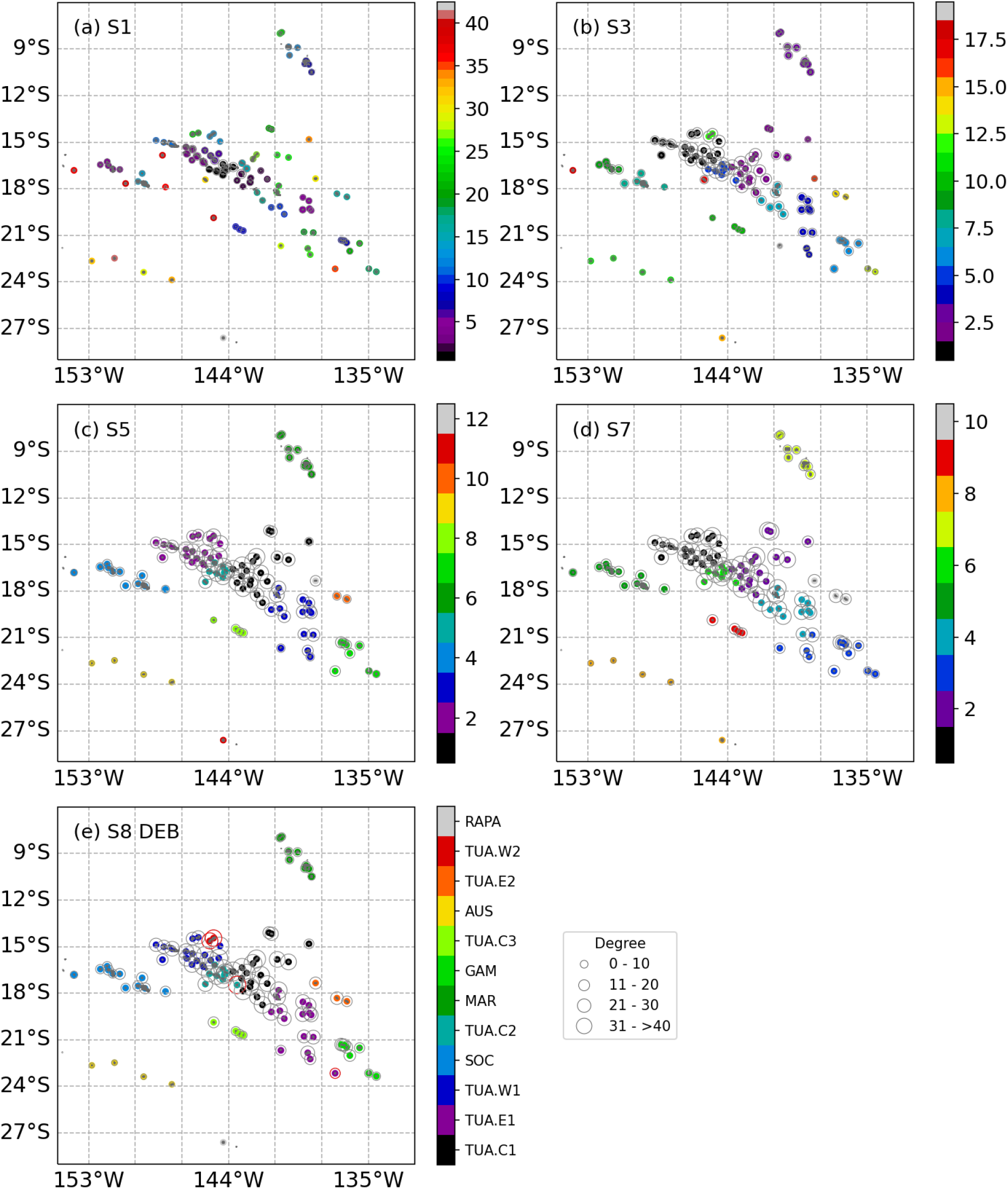
Cluster distributions over the 10-year simulation dataset for (a) scenario 1, (b) scenario 3, (c) scenario 5 and (d) scenario 7 and (e) scenario 8 (DEB). Colors refer to clusters sorted from the largest to the smallest one from bottom to top. Grey circles refer to the number of degree. In (e), circles in red represent the islands that changed clusters between scenario 5 and scenario 8.

### 3.4 *Pinctada margaritifera* connectivity (Scenario 8)

At FP scale, the LPP obtained from the DEB model increased southward, with a minimum of 11.55 days for Marquesas islands and reaching >27-30 days for Austral islands, Gambier islands, and the southeastern part of Tuamotu archipelago (Table 3). Both seasonal and interannual variations of SS for *P. margaritifera* were stronger than for the fixed LPP scenarios (Figure 3). The potential connectivity matrix is comparable to the one from scenario 7 with a fixed LPP of 23 days (Figure 4d and 4e). However, by considering environmental influences on LPP, potential connectivity increases within the Marquesas and western Tuamotu archipelagos (2-5%) while it generally decreases in the others (Figure 4f). Network clustering based on connectivity matrices computed using the DEB model over the 10 years simulated provided 12 different clusters (Figure 6e). The Society (SOC), Gambier (GAM) and Marquesas (MAR) archipelagos were well delimited. Austral (AUS) archipelago was identified as a separate cluster, but its remotest island Rapa formed a cluster on its own while the Tuamotu archipelago was divided in 7 clusters. These patterns are close to those observed in scenario 5 with a fixed LPP of 17 days (Figure 6c); however, some differences were observed among the islands. Some of them formed new clusters on their own in scenario 8 (DEB) such as Takaroa and Takapoto in the western Tuamotu. While the others changed from one to another in the eastern Tuamotu (Morane) and in the central Tuamotu (Haraiki). Average SS varied across clusters (K-W Chi-squared = 57.7, *P* < 0.001), ranging from 0.18% ± 0.20 to 30.61% ± 4.27 for RAPA and TUA.W2, respectively (Table 3). Similarly, RS varied largely across clusters (K-W Chi-squared = 114.88, *P* < 0.001), with again an almost null RS for RAPA, while maximum RS was observed for TUA.W1 (Table 3). SR varied across clusters (K-W Chi squared = 109.78, *P* < 0.001), reaching 100% for AUS, RAPA and MAR in scenario 8 (DEB). LR was higher in the West Tuamotu (TUA.W1 and TUA.W2) reaching 16.74 ± 1.67 and 14.10 ± 6.53, respectively (Table 3). TUA.C2 showed the highest degree value, indicative of the centrality of the cluster for the network (Table 3).

**Table 3:** Summary statistics for theoretical dispersal based on DEB model for P. margaritifera (scenario 8) and 10 years computation (2010-2019). Letters represent significant differences across clusters at Bonferroni adjusted P < 0.05.

| Clusters | SS (%) | SR (%) | LR (%) | RS (%) | Degree | Distance to settlement (km) | LPP (days) |
| --- | --- | --- | --- | --- | --- | --- | --- |
| AUS | $0.43 \pm 0.27^a$ | $100 \pm 0.00^a$ | $0.43 \pm 0.27^a$ | $0.01 \pm 0.01^a$ | 3.50 | $96.15 \pm 50.03^{abc}$ | $29.5 \pm 6.60^{abcd}$ |
| GAM | $1.77 \pm 0.44^b$ | $96.85 \pm 4.35^{bcde}$ | $1.64 \pm 0.45^b$ | $0.13 \pm 0.04^b$ | 15.50 | $92.24 \pm 30.04^{abc}$ | $28.0 \pm 1.25^a$ |
| MAR | $3.73 \pm 1.40^c$ | $100.00 \pm 0.00^{ab}$ | $3.73 \pm 1.40^c$ | $0.33 \pm 0.12^c$ | 8.67 | $50.45 \pm 7.74^a$ | $11.5 \pm 1.42^e$ |
| RAPA | $0.18 \pm 0.20^a$ | $100.00 \pm 0.00^{ab}$ | $0.12 \pm 0.20^a$ | $0.00 \pm 0.00^d$ | 1 | $5.77 \pm 3.23$ | $30.0 \pm 12.89^{abcde}$ |
| SOC | $7.00 \pm 1.04^{de}$ | $98.13 \pm 2.11^c$ | $6.99 \pm 1.03^{de}$ | $0.77 \pm 0.11^e$ | 12.45 | $73.62 \pm 11.96^b$ | $22.5 \pm 0.96^{bc}$ |
| TUA.C1 | $9.31 \pm 1.39^d$ | $83.07 \pm 7.86^f$ | $6.59 \pm 1.14^{de}$ | $1.41 \pm 0.27^f$ | 33.55 | $120.63 \pm 8.89^c$ | $23.5 \pm 2.81^{bcd}$ |
| TUA.C2 | $14.70 \pm 2.84^f$ | $61.75 \pm 6.52^g$ | $9.43 \pm 1.74^{df}$ | $1.49 \pm 0.19^f$ | 36.40 | $76.81 \pm 13.46^b$ | $23.5 \pm 2.13^{bc}$ |
| TUA.C3 | $0.67 \pm 0.47^a$ | $88.03 \pm 8.73^{cdef}$ | $0.66 \pm 0.46^a$ | $0.03 \pm 0.02^a$ | 10.75 | $59.39 \pm 32.92^{ab}$ | $30.0 \pm 2.62^{ad}$ |
| TUA.E1 | $3.49 \pm 1.54^{bc}$ | $79.11 \pm 14.62^{dfg}$ | $3.09 \pm 1.34^{bc}$ | $0.60 \pm 0.20^{ce}$ | 24.68 | $100.68 \pm 18.13^{bc}$ | $27.0 \pm 2.51^{abd}$ |
| TUA.E2 | $4.30 \pm 2.64^{ce}$ | $97.97 \pm 2.69^{abce}$ | $4.39 \pm 2.55^{bce}$ | $0.13 \pm 0.07^b$ | 17.33 | $55.82 \pm 28.91^{abc}$ | $24.5 \pm 2.46^{abcd}$ |
| TUA.W1 | $19.16 \pm 2.48^f$ | $76.87 \pm 3.57^d$ | $16.74 \pm 1.67^g$ | $3.42 \pm 0.37^g$ | 27.81 | $88.12 \pm 7.72^b$ | $20.5 \pm 2.50^c$ |
| TUA.W2 | $30.61 \pm 4.27^g$ | $86.05 \pm 7.22^{def}$ | $14.10 \pm 6.53^{fg}$ | $0.32 \pm 0.13^c$ | 33.00 | $90.77 \pm 27.41^{bc}$ | $20.0 \pm 2.72^c$ |

## 4 Discussion

### Connectivity in FP and consequences for management

Ocean connectivity is one of the main drivers of metapopulation dynamics and evolution in marine organisms. Larval transport and dispersal potential are constrained both by i) hydro-physical properties of the environment, notably current transport, and by ii) genetic and phenotypically plastic properties of the organism which determine LPP and PLD. To assess the relative role of LPP on FP connectivity we used a Lagrangian particle dispersal model through 8 scenarios that considered various LPP and maximum PLD (from 5 to 35 days). Using a graph-theory framework (Treml et al., 2008), we showed that population fragmentation, estimated through overall network density and the number of identified clusters, decreased with increasing LPP and maximum PLD. The association between PLD and population connectivity has also been detected using population genetic approaches, notably based on isolation-by-distance (IBD) estimations derived from genetic data (Selkoe & Toonen, 2011; Treml et al., 2012). As PLD varies substantially in the marine realm, from 0 for lecitrophic to >100 days for some planktonic species (Mercier et al., 2013), our observation that the clustering of populations in FP depends on PLD highlights the importance of defining species-specific management units for fishery management and for live animal translocation for aquaculture purposes. Furthermore, we showed that FP archipelagos remain largely disconnected, even for the 35-days-maximum-PLD scenario, with 12 main clusters identified. Such isolation was also observed in other species like giant clams (*Tridacna maxima;*<u>Laurent et al., 2002)</u>, whose LPPs are shorter than those of *P. margaritifera*.

### Environmental drivers of connectivity

Among the environmental conditions that could affect settlement rates and PLD, temperature has been one of the most studied in marine species (O’Connor et al., 2007; Raventos et al., 2021), and little attention has been given to the influence of food availability. Nevertheless, in FP, temperatures are relatively stable compared to temperate regions, suggesting that this might not be the main driver influencing PLD and connectivity (Sangare et al., 2020a). In turn, Chl-a concentration exhibits greater variability across the territory. Chla is significantly higher in the Marquesas than at any other archipelago due to a strong island mass effect, while temperature remains close to the optimal for *P. margaritifera* (Le Moullac et al., 2016; Raapoto et al., 2018, 2019). According to empirical observation and modeling for *P. margaritifera* (Sangare et al., 2020a; Thomas et al., 2011), these environmental conditions decrease LPP and favor local retention of larvae, as has been previously documented (Raapoto et al., 2018). Our results support the observation of a segregation between the Marquesas and the rest of FP based on microsatellite data (Reisser et al., 2019).

Overall, our results suggest that the contribution of the environmental conditions to *P. margaritifera* clustering at the scale of FP is dampened, and connectivity is mainly driven by the geographical configuration of the archipelagos. The Marquesas archipelago (in the North) is isolated by both distance and short LPP due to Chl-a rich waters. The Austral archipelago (in the South) remains isolated from the rest of the network mainly because of geographical and hydrodynamical reasons. Even if temperature experienced by larvae in the area tend to extend their LPP, the main current flow and the remoteness of these islands prevent connectivity with other northern clusters. Finally, the islands of the Society and Tuamotu archipelagos form a denser group with homogeneous environmental conditions which lead to clustering patterns consistent with what is observed with the fixed 23-days-LPP scenarios. Our analysis also revealed that, even though DEB model predictions do not enhance estimates of connectivity among highly-fragmented clusters, they do provide valuable information when considering dispersal metrics at narrower temporal and spatial scales. The variability in temperature and Chl-a modulates the settlement success at seasonal and interannual scales (Fig. 3), and can also sensibly identified clusters, such as Takaroa and Takapoto in the western Tuamotu (Fig. 6e), Takapoto being the main provider of spat for the pearl farming sector.

### Consequence for management

Results from our model can inform science-based management practices. The pearl industry in FP depends on wild spat collection. A few atolls (5 to 6) provide most of the biological material to the 556 pearl producers across 26 atolls and islands (Ky et al. 2019). This system has favored the spread of parasites and diseases and led to the regulation of transfers [i.e., destination restricted to non-collecting atolls and limited to locally collected biological material in the Gambier archipelagos, (Bondad-Reantaso et al., 2007; Chagot et al., 1993) and genetic homogenization (Lemer & Planes, 2012). We show that spat-collecting atolls, which are mainly located in the West Tuamotu, are the most central in the network, with elevated connectivity. Moreover, atoll connectivity help maintaining stable populations (Hughes, 1990), hence, informing on population resilience potential. Such information is vital for tropical ectotherm species leaving already close to their upper thermal limits (Hughes et al., 2003; Seebacher et al., 2014), and for enclosed lagoons which are sensitive to environmental degradation (Andréfouët et al., 2015; Rodier et al., 2019).

### Relevance of the approach and results for other species and archipelagic countries

The dispersal modelling approach developed in this study was justified for *P. margaritifera*‘s case study in FP, but might require adjustments if used for other species or archipelagos. First, the currents data (GLORYS) we used to force the Lagrangian model had a relatively coarse spatial and temporal resolution (∼9 km; daily). Although FP islands are located in a low eddy-activity area (Chelton et al., 2007), the presence of obstacles in the flow field may induce turbulent flow downstream. Currents used here do not resolve sub-mesoscale dynamics around islands, which would allow to determine the proximity of particles to passes and their potential influx towards the lagoon. The low resolution of currents used in this study was of limited concern for characterizing the general patterns of connectivity at the scale of the territory, considering the wide distance between many islands in FP. Also, current velocities being averaged across the first 50 m depth, Stokes drift forcing is not considered in our experiment since its magnitude decrease rapidly with depth (Tamura et al., 2012). Implementing a similar approach in a less fragmented system, or at smaller scale, by contrast, may require adapting the resolution of the model to picture adequately the sub-mesoscale processes.

Second, in our model, particles were limited to a 2-dimensional displacement only. This was justified for *P. margaritifera* whose larvae distribution is light dependent so that larvae can maintain themselves in the upper layers of the water column (Thomas, Garen, et al., 2012), but this may be a limit to transpose results to species whose larvae occupy the water column differently.

Third, the dispersal kernels modelled in this study do not account for differences in population abundance between islands. Differences in populations abundance among islands do not change the probability of transport from an island i to an island j (*P_i→j_*), nor local retention (LR), but may change estimates for other dispersal metrics calculated here (SS, RS, and SR), as well as the degree and number of clusters derived from graph theory. To date, *P. margaritifera* natural and farmed stocks have been assessed only for a few lagoons in FP (Andréfouët et al., 2016; Bionaz et al., 2022), which precludes us from integrating this variable into the dispersal models. While population size could be related with atoll size in other regions, this is not the case in FP (Andréfouët et al., 2016; Bionaz et al., 2022). Full estimates of *P. margaritifera* stocks would allow to weight larval emission by localization, and potentially improve the connectivity matrices (Falcini et al., 2020). Similar improvement would benefit from the inclusion of additional forces modulating effective connectivity (e.g., spawning, and larval and post-larval survival). Meanwhile, one can hypothesize that the pattern of connectivity estimated in this study (i.e., high isolation of populations in Marquesas and Austral archipelagos compared to other archipelagos) remain probably true, and is even more accentuated in reality.

Fourth, the percentage of larval export from the lagoon to oceanic waters might be linked to lagoon water renewal, which vary depending on the atoll’s geomorphology (Andréfouët et al., 2022). In the semi-enclosed atoll lagoon of Ahe, larval export rate has been estimated to be ∼40% (Andréfouët et al., 2021b), while closed atolls are expected to export significantly less larvae (Andréfouët et al., 2001). Although the dispersal kernels we estimated do not consider the degree of openness of each island/atoll, our results provide a basis of dispersal patterns triggered by oceanic transport at the scale of the territory. This can contribute to finer-grain dispersal studies that will take local geomorphology and mesoscale circulation patterns into account.

Our model integrating spatial-DEB (scenario 8) identified 12 clusters, which largely overlapped with existing archipelago-wise management units, and is congruent with previous genetic data for *P. margaritifera*. Specifically, our simulations clearly identified Gambier, Marquesas and Austral islands as disconnected from the Tuamotu Archipelago, in accordance with previous genetic clustering studies for *P. margaritifera* (Lemer & Planes, 2014; Reisser et al., 2019). West Tuamotu atolls were at the center of the network showing the highest connectivity with surrounding atolls and islands. The archipelago itself was divided into 7 different clusters, but this was not detected with genetic data (Lemer & Planes, 2014; Reisser et al., 2019). The discrepancy is likely due to both the confounding effect of human transfers, and the different scales of observation used in our model and the population genetic approach (long time-scale observations *versus* single-step connectivity). Models accounting for dispersion with steppingstones and filial-coalescence connectivity, improved knowledge on spawning, and stock assessments would certainly help in resolving connectivity in the Tuamotu (Boulanger et al., 2020; Legrand et al., 2022).

### Impact of climate change on larval dispersal

By examining the larval dispersal and connectivity patterns of different marine bivalve species including *P. margaritifera*, within FP islands on the 2010-2019 period, we found variable settlement success (SS) depending on ENSO cycles. Notably, lower SS were found during strong La Niña periods. This result raises the question of the potential impact of climate change on connectivity patterns in FP. Ocean warming is widely considered to be one of the major stressors to ocean ecosystems <u>(Raventos et al., 2021)</u>. Rising temperature has a direct impact on larval development by decreasing PLD. At global scale, a PLD decrease of 10% to 25% is suspected . Such decrease might isolate the Marquesas and Austral archipelago even more. Furthermore, this change in temperature may affect the timing of reproduction in many marine species, including *P. margaritifera.* The time lag induced by the temperature change might modify the connectivity and its variability for certain species (Lacroix et al., 2018). Furthermore, the frequency, intensity, and duration of marine heat waves is expected to increase in the future (Rahmstorf & Coumou, 2011). Similar approaches using DEB modeling but using ocean projection instead of hindcast data is the next to better capture the potential impact of these warming events on connectivity at regional scale. Given the homogeneous temperature pattern in FP, such perspective work should not only consider ocean temperature projection, but also the change in ocean circulation in response to climate change. Indeed, these changes in ocean conditions could also alter larval transport pathways, potentially affecting the dispersal patterns and genetic diversity of many species. Previous studies suggest that the change in ocean circulation might be the most prominent driver of the dispersal in a climatechange context (Cetina-Heredia et al., 2015). Finally, climate change is likely to lead to a decrease in global phytoplankton biomass (Boyce et al., 2010). This decrease in phytoplankton could have significant repercussions on food availability for larvae development and thus extending their LPP. Therefore, it is important that future research continues to investigate the impact of climate change on larval dispersal and the implications for the genetic diversity and sustainability of marine species.

## 5 Conclusions

This paper assessed the role of pelagic larval duration and population connectivity in marine organisms, with a focus on the black-lip pearl oyster *P. margaritifera* in FP. We used a Lagrangian particle dispersal model to recreate seven scenarios of maximum PLD ranging from 5 to 35 days and one scenario based on a DEB modeling approach. The graph-theory framework showed that even though these archipelagos are not fully connected, population fragmentation decreases with increasing PLD.

The wealth of biological data available for the local model species *P. margaritifera* allowed producing a biophysical dispersion model that explicitly consideres the energetic condition of individual larvae, and predict settlement success. Even though the DEB model derived predictions did not modify the regional clustering in FP, it did help explain the interannual and seasonal variability of settlement success. These results further highlight the importance of accounting for variability in food and temperature for assessing the impacts of climate change over the regional connectivity.

The results indicate the importance of defining species-specific management units for fishery management and for live animal translocation for aquaculture purposes in FP and other archipelagic countries. Environmental drivers such as temperature and food availability were also discussed, with a focus on the influence of Chl-a concentration on local retention of larvae. Our approach lends itself for investigating the connectivity patterns of other species or regions, thus providing a valuable tool to improve our understanding of marine metapopulation dynamics and evolutionary processes.

## Data Availability

The data that supports the findings of this study and used to train the given model are available from the corresponding author upon reasonable request.

## Acknowledgments

The authors acknowledge the Pôle de Calcul et de Données Marines (PCDM; https://wwz.ifremer.fr/en/Research-Technology/Research-Infrastructures/Digital-infrastructures/Computation-Centre) for providing DATARMOR computing and storage resources.

## Author contributions

JLL, CJM, SVW and R.L conceived the project and collected the data. HR and RL collected the data and built the physical model. CJM and HR collected the data and run the DEB model. HR, SVW and JLL conducted the analyses of the dispersal data. HR and JLL drafted the manuscript. All authors contributed to the writing and have reviewed the manuscript.

## Supplementary Materials

**S1 tab:** Indexes of islands assigned following the main current with the name of the cluster to which it belongs and, the average LPP.

| Index | Island | Cluster | LPP | Index | Island | Cluster | LPP | Index | Island | Cluster | LPP |
| --- | --- | --- | --- | --- | --- | --- | --- | --- | --- | --- | --- |
| 1 | Hatutu | MAR | 12.06 | 35 | Tuanake | TUA3 | 23.11 | 69 | Maiao | SOC | 22.90 |
| 2 | Eiao | MAR | 11.90 | 36 | Motutunga | TUA3 | 21.85 | 70 | Huahine | SOC | 21.04 |
| 3 | NukuHiva | MAR | 12.80 | 37 | Tahanea | TUA3 | 22.54 | 71 | RaiateaTahaa | SOC | 22.33 |
| 4 | UaHuka | MAR | 12.55 | 38 | Faaite | TUA3 | 21.88 | 72 | BoraBora | SOC | 21.36 |
| 5 | UaPou | MAR | 13.15 | 39 | Anaa | TUA3 | 23.22 | 73 | Tupai | SOC | 22.11 |
| 6 | HivaOa | MAR | 13.75 | 40 | Katiu | TUA3 | 22.71 | 74 | Maupiti | SOC | 22.47 |
| 7 | Mohotane | MAR | 12.34 | 41 | Raraka | TUA3 | 22.40 | 75 | Maupihaa | SOC | 24.53 |
| 8 | Tahuata | MAR | 13.11 | 42 | Taiaro | TUA1 | 21.27 | 76 | PukaPuka | TUA2 | 19.56 |
| 9 | FatuHiva | MAR | 12.12 | 43 | Kauehi | TUA3 | 20.22 | 77 | Tatakoto | TUA2 | 26.49 |
| 10 | Tepoto | TUA2 | 19.97 | 44 | Aratika | TUA1 | 21.11 | 78 | Pukarua | TUA2 | 23.72 |
| 11 | Napuka | TUA2 | 19.88 | 45 | Fakarava | TUA3 | 20.28 | 79 | Reao | TUA2 | 23.06 |
| 12 | Fangatau | TUA2 | 20.88 | 46 | Toau | TUA1 | 19.73 | 80 | Akiaki | TUA2 | 25.80 |
| 13 | Fakahina | TUA2 | 22.07 | 47 | Niau | TUA1 | 20.59 | 81 | Vahitahi | TUA2 | 26.38 |
| 14 | Takume | TUA2 | 21.76 | 48 | Kaukura | TUA1 | 20.64 | 82 | Nukutavake | TUA2 | 27.26 |
| 15 | Raroia | TUA2 | 22.67 | 49 | Apataki | TUA1 | 18.06 | 83 | Pinaki | TUA2 | 27.51 |
| 16 | Taenga | TUA2 | 23.31 | 50 | Arutua | TUA1 | 17.82 | 84 | Vairaatea | TUA2 | 26.03 |
| 17 | Nihiru | TUA2 | 24.13 | 51 | Rangiroa | TUA1 | 20.51 | 85 | Vanavana | TUA2 | 29.03 |
| 18 | Rekareka | TUA2 | 23.16 | 52 | Tikehau | TUA1 | 21.55 | 86 | Tureia | TUA2 | 22.78 |
| 19 | Tauere | TUA2 | 23.81 | 53 | Mataiva | TUA1 | 21.02 | 87 | Moruroa | GAM | 27.22 |
| 20 | Amanu | TUA2 | 25.69 | 54 | Makatea | TUA1 | 22.62 | 88 | Fangataufa | GAM | 27.46 |
| 21 | Hao | TUA2 | 24.22 | 55 | Takaroa | TUA1 | 20.45 | 89 | Tenararo | GAM | 29.12 |
| 22 | Paraoa | TUA2 | 25.90 | 56 | Takapoto | TUA1 | 20.50 | 90 | Vahanga | GAM | 29.74 |
| 23 | Ahunui | TUA2 | 26.27 | 57 | Tikei | TUA1 | 21.34 | 91 | Tenarunga | GAM | 30.12 |
| 24 | Manuhangi | TUA2 | 26.41 | 58 | Manihi | TUA1 | 21.51 | 92 | Matureivavao | GAM | 29.27 |
| 25 | NengoNengo | TUA2 | 25.86 | 59 | Ahe | TUA1 | 20.26 | 93 | MaruteaSud | GAM | 28.44 |
| 26 | RavahereMarokau | TUA2 | 25.28 | 60 | Anuanuraro | GAM | 27.56 | 94 | Maria | GAM | 30.94 |
| 27 | Reitoru | TUA2 | 25.19 | 61 | Anuanurunga | GAM | 28.65 | 95 | Mangareva | GAM | 21.77 |
| 28 | Hikueru | TUA2 | 23.23 | 62 | Nukutepipi | GAM | 27.95 | 96 | Temoe | GAM | 26.20 |
| 29 | Tekokota | TUA2 | 24.02 | 63 | Hereheretue | GAM | 24.59 | 97 | Morane | GAM | 30.93 |
| 30 | Haraiki | TUA3 | 23.74 | 64 | Tematangi | GAM | 27.72 | 98 | Rimatara | AUS1 | 30.78 |
| 31 | MaruteaNord | TUA2 | 23.95 | 65 | Mehetia | SOC | 23.38 | 99 | Rurutu | AUS1 | 29.31 |
| 32 | Makemo | TUA2 | 23.38 | 66 | Tetiaroa | SOC | 22.00 | 100 | Tubuai | AUS1 | 27.14 |
| 33 | TepotoSud | TUA3 | 22.26 | 67 | Tahiti | SOC | 23.15 | 101 | Raivavae | AUS1 | 30.41 |
| 34 | Hiti | TUA3 | 23.83 | 68 | Moorea | SOC | 22.70 | 102 | Rapa | AUS2 | 28.22 |

\*FILE ANIMATION

*FILE ANIMATION

*S2 File. Animation of the modelled dispersal paths for a duration of 35 days without settlement on FP, expressed as larval densities and calculated as the number of particles in each cell on a 1/100° grid*.

**S3 Fig.**
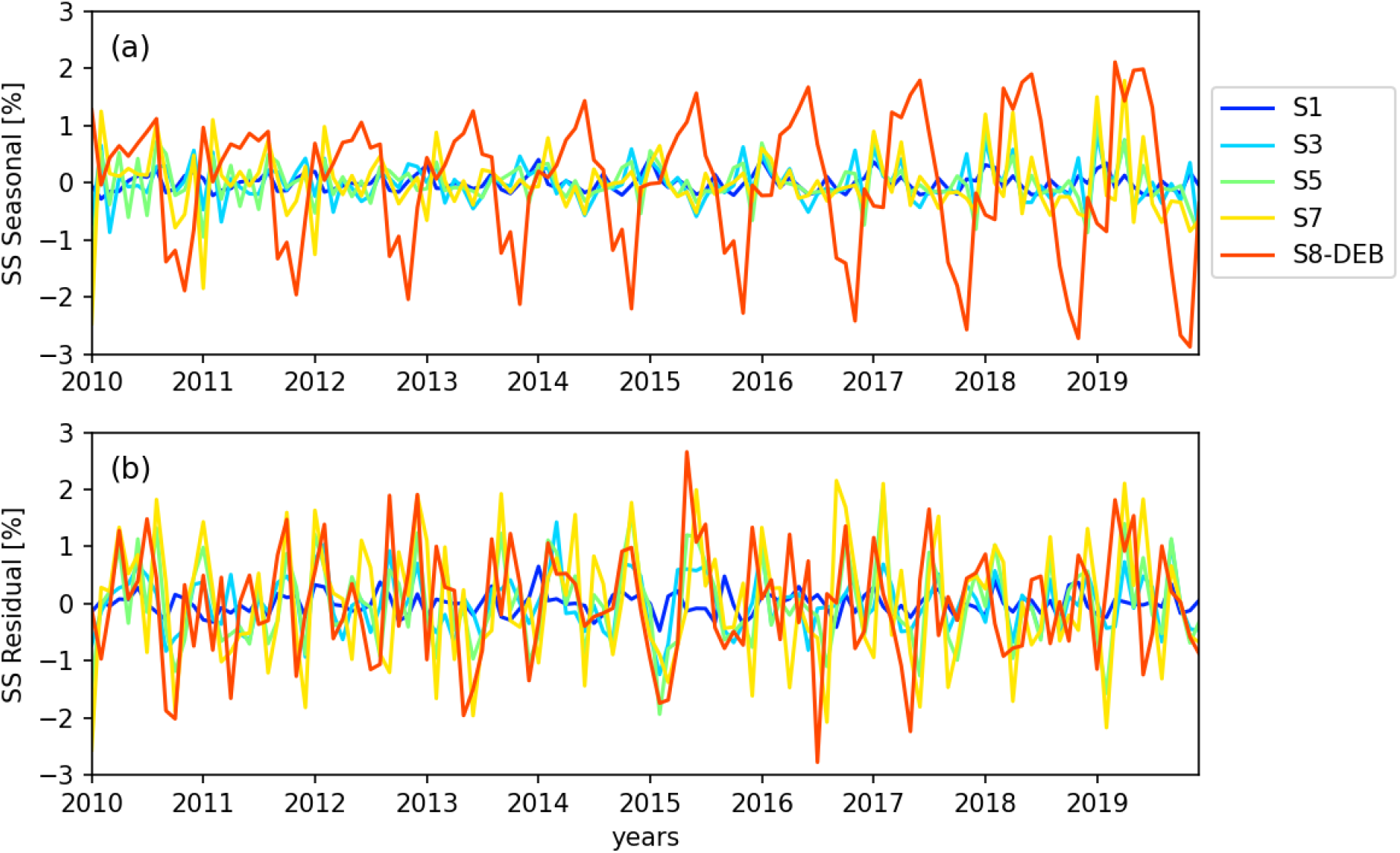
(a) Seasonal and (b) residual settlement success (SS) extracted from the STL over the full dataset (all archipelagos included). For ease of representation, only scenarios 1, 3, 5, 7 and 8 are represented in blue, light blue, green, yellow and orange, respectively.

