## Supplementary figures and images for "Can food and temperature influence regional connectivity patterns of Bivalvia in fragmented archipelagos? Evidence from biophysical modeling applied to French Polynesia"

### FigureS2_Raapoto_etal_bioRxiv.gif

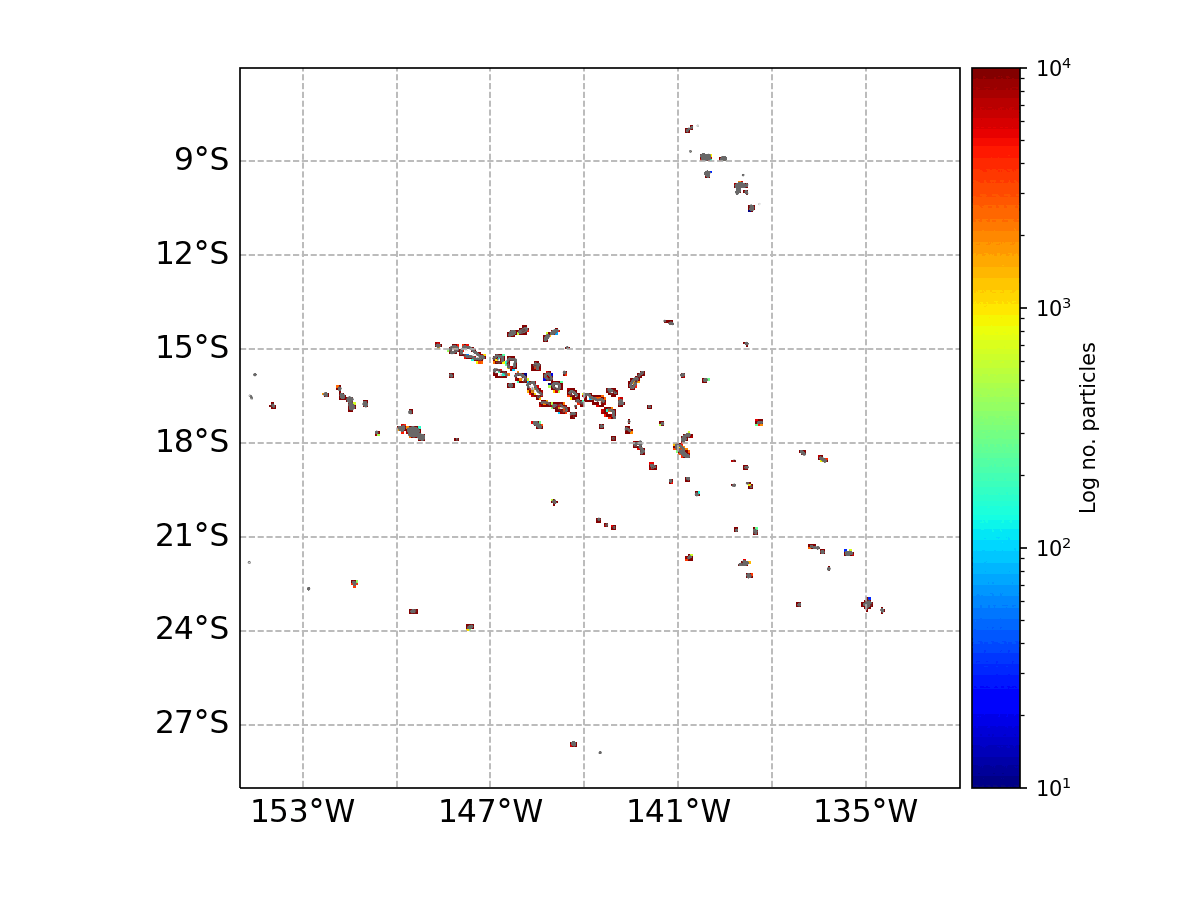
